# Lateralization of autonomic activity in response to limb-specific threat

**DOI:** 10.1101/2021.11.24.469931

**Authors:** James H. Kryklywy, Amy Lu, Kevin H. Roberts, Matt Rowan, Rebecca M. Todd

## Abstract

In times of stress or danger, the autonomic nervous system (ANS) signals the *fight* or *flight* response. A canonical function of ANS activity is to globally mobilize metabolic resources, preparing the organism to respond to threat. Yet a body of research has demonstrated that, rather than displaying a homogenous pattern across the body, autonomic responses to arousing events — as measured through changes in electrodermal activity (EDA) — can differ between right and left body locations. Surprisingly, the metabolic function of such ANS asymmetry has not been investigated. In the current study, we investigated whether asymmetric autonomic responses could be induced through limb-specific aversive stimulation. Participants were given mild electric stimulation to either the left or right arm while EDA was monitored bilaterally. Across participants, a strong ipsilateral EDA response bias was observed, with increased EDA response in the hand adjacent to the stimulation. This effect was observable in over 50% of individual subjects. These results demonstrate that autonomic output is more complex than canonical interpretations suggest. We suggest that, in stressful situations, autonomic outputs can prepare either the whole-body *fight* or *flight* response, or a simply a limb-localized *flick*, which can effectively neutralize the threat while minimizing global resource consumption. These findings provide insight into the evolutionary pathway of neural systems processing general arousal by linking observed asymmetry in the peripheral arousal response to a historical leveraging of neural structures organized to mediate responses to localized threat.

## Introduction

An organism’s ability to respond efficiently to threatening situations can mean the difference between survival and death. In the presence of an acute stressor, the autonomic nervous system (ANS) - specifically, the sympathetic nervous system – signals the body to prepare for action. Such ANS activation results in increases in cardiac and respiratory outputs, dilation of vasculature in skeletal muscles (in mammals), release of glucocorticoids into the bloodstream, and increased electrodermal activity (EDA; a measure of sweat gland permeability as observed through resistance to small electrical currents; Boucesein, 2012; Critchley, 2002). This ‘fight or flight’ response is highly conserved across vertebrate (Romero, 2019; Romero and Wingfield, 2016) and some invertebrate (Shimizu and Okabe, 2007) species and its effective manifestation is critical to the deployment of specific survival behaviors (Barrios et al., 2021; Blas et al., 2007; Lee and Wang, 2019; Romero, 2019).

The canonical role of the ANS is the mobilization or conservation of metabolic resources as demanded by situational cueing (Brener, 1987; Gibbons, 2019; Wehrwein et al., 2016). Emotional arousal in response to relevant events can trigger a sympathetic nervous response (for extended review, see Kreibig, 2010) including increases in EDA (Vetrugno et al., 2003). Early theories proposed that such responses are homogenous across the entire body (Cannon, 1939). Although the field has moved on, current working assumptions still maintain generalized output to, and responses from, individual *effector organs* (Folkow, 2000; Janig and McLachlan, 1992) mediated by a centralized autonomic network (Benarroch, 1993; Damasio, 1998; Valenza et al., 2019). Notably, with respect to the electrodermal-effector organ, the contemporary division of ANS function still preserves the idea that the skin receives homogenous sympathetic output signaling the need for motor preparedness (Blakemore and Vuilleumier, 2017; Fredrikson et al., 1998; Le et al., 2019). Yet there is a substantial body of research (e.g., Bjorhei et al., 2019; i Baque et al., 1984; Picard et al., 2016; Richter, 1927) demonstrating that asymmetric ANS responses — as measured by changes in electrodermal activity (EDA) — can differ between right and left body locations. One potential function of such asymmetry could be a limb-specific response to threat directed to one side of the body, challenging the assumption of global homogeneity. Surprisingly, the question of whether asymmetric ANS responses result from limb-specific threat has been almost wholly neglected.

Historical data from as early as the 1920’s (Richter, 1927; Syz, 1926) provides evidence that ANS outputs – specifically those monitored through EDA - are not always consistent across the body. Examination and explanation of these effects throughout the first half of the 20^th^ century were minimal, relying predominantly on pathological (Fisher, 1958; Fisher and Cleveland, 1959; Galbrecht et al., 1965) or structural (i.e., lesion dependent, Holloway and Parsons, 1969; Richter, 1927) justification for asymmetry, albeit with some theories pairing EDA asymmetry to degrees of general arousal (Obrist, 1963). While interest in the area of lateralized EDA in both neuro-typical and atypical populations intensified in the 1970’s (for review, see i Baque et al., 1984), this was paired to the rise of theories evoking gross hemispheric specialization (i.e., right brain vs. left brain rhetoric Galaburda et al., 1978; Reeves, 1983; Sperry, 1968), and overlooked what minimal evidence existed for a physiological basis for asymmetric ANS architecture (Fuhrer, 1971). As a consequence, much of this work fell into disrepute alongside the repudiation of the overarching frameworks in which they were nested.

Recently, incidental findings from the field of computer science have reinvigorated interest in EDA asymmetry. In data from wearable electrophysiological recording devices (Ayzenberg and Picard, 2014; Poh et al., 2012; Poh et al., 2010; Sano et al., 2014), collected for the purpose of training computer algorithms to sense, recognize and respond to human emotional information (e.g., el Kaliouby et al., 2006; Picard, 2002; Picard, 2009), asymmetric EDA activity has been observed in response to specific types of emotional situation or arousal (Picard et al., 2016). This work has prompted a secondary resurgence of study on the lateralization of ANS outputs (e.g., Banks et al., 2012; Bjorhei et al., 2019; Kasos et al., 2018; Picard et al., 2016) primarily focused on understanding how data from wearable devices can be used to index biomarkers for mental health monitoring (Arza et al., 2019; Ghandeharioun et al., 2017; Greene et al., 2016; Mohr et al., 2017) and clinical impairment (e.g., autism, Baker et al., 2018; addiction, Carreiro et al., 2015a; Carreiro et al., 2015b; dementia, Kourtis et al., 2019). Stemming from this line of research, the *multiple sources of arousal* theory (Picard, 2015) points to evidence that asymmetric EDA activation can result from *ipsilateral* signals from “limbic” regions, in particular the amygdalae, linked to stress or emotional arousal. It can also arise from *contralateral* signals from basal ganglia and premotor regions linked to motor preparedness.

In the context of canonical views of ANS functioning, however, – ANS activation as a mechanism to mediate metabolic resource allocation – there is no clear reason why centrally-mediated arousal requiring whole body-responses would elicit greater EDA activity lateralized to one side versus another. Yet, the underlying architecture of the autonomic nervous system is such that left-right signal variability undeniably occurs (Banks et al., 2012; Bjorhei et al., 2019; i Baque et al., 1984; Obrist, 1963; Picard et al., 2016; Richter, 1927). Interestingly, however, almost all previous work displaying asymmetric EDA has involved stimuli that elicit generalized states of arousal not requiring body-part specific responding (e.g., face perception, Banks et al., 2012; high vs. low impact stressors, Bjorhei et al., 2019; situational arousal, i Baque and de Bonis, 1983; Obrist, 1963; Picard et al., 2016). However, it is unclear how asymmetric responding to generalized arousal could serve a functional purpose within the canonical role of the ANS – that of metabolic conservation - and thus it is unlikely that these stimuli are the primary developmental motivators of the observed lateralized architecture. Rather, centrally mediated arousal processing is likely leveraging neural architecture based on physical responses to direct threat, where lateralization of response matters. Notably, there is almost no reference made to research conducted on direct tangible limb-specific tactile threat. Yet one largely forgotten historical assay provides preliminary support for the hypothesis that responses to limb-specific threat are lateralized (Fuhrer, 1971). Such preliminary evidence supports the hypothesis that a localized proximal threat to an organism may not require whole body action, but rather a targeted response in the threatened limb. Maximal conservation of metabolic resources would occur if sympathetic activation is evoked in the specific limbs required for the motion that will allow a return to safety, while the rest of the body maintains a state of rest.

The aim of the present study was to investigate lateralized changes in EDA in response to limb-specific aversive stimulation. Specifically, we aimed to identify whether increased sympathetic output is directed to a threatened limb, indicating a potential increase in resource mobilization limited to the site requiring subsequent motor response. To assess lateralization biases in EDA responses, we compared EDA activity from each arm during ipsi- and contra-lateral stimulation. Consistent with theories of metabolic conservation in the autonomic nervous system (Gibbons, 2019; Wehrwein et al., 2016), we predicted that EDA responses would be greater during ipsilateral stimulation, and that this bias would be observable when recording from both the left and right arm.

## Results

To assess lateralization biases, we contrasted electrodermal activity (EDA) between the left and right hands in response to mild electric stimulation applied either ipsi- or contra-laterally to the galvanic skin response (GSR) electrodes. To obtain an index of ‘Lateralization Bias’, z-score standardized GSR data collected from the left hand was subtracted from that collected from the right hand. This data was then split into three distinct trial types: Left shock, Right Shock, and No Shock. All presented *p* statistics have been subject to a false-detection rate correction (Benjamini and Hochberg, 1995).

### Across-subject analyses

To assess lateralization biases across all participants, a series of one-sample time-locked t-tests were conducted comparing the observed Lateralization Bias against a test value of zero (i.e., no difference in left versus right side EDA response) for each Left-shock, Right shock and No Shock conditions averaged within participant. This identified a clear lateralized response bias in both left and right recording sites for ipsilateral stimulation (Left Shock: onset = 1920 ms, offset = 4840 ms, |t_range(49)_| = 2.69-5.68, all *p* < 0.05; Right Shock: onset = 2790 ms, offset = 5490 ms, |t_range(49)_| = 2.68-6.93, all *p* < 0.05; Figure 1): Greater EDA was observed at the recording site on the same side as the shock was administered relative to the contralateral recording site. By contrast, Lateralization Bias did not differ from zero at any time point during ‘No Shock’ trials.

**Figure 1.**
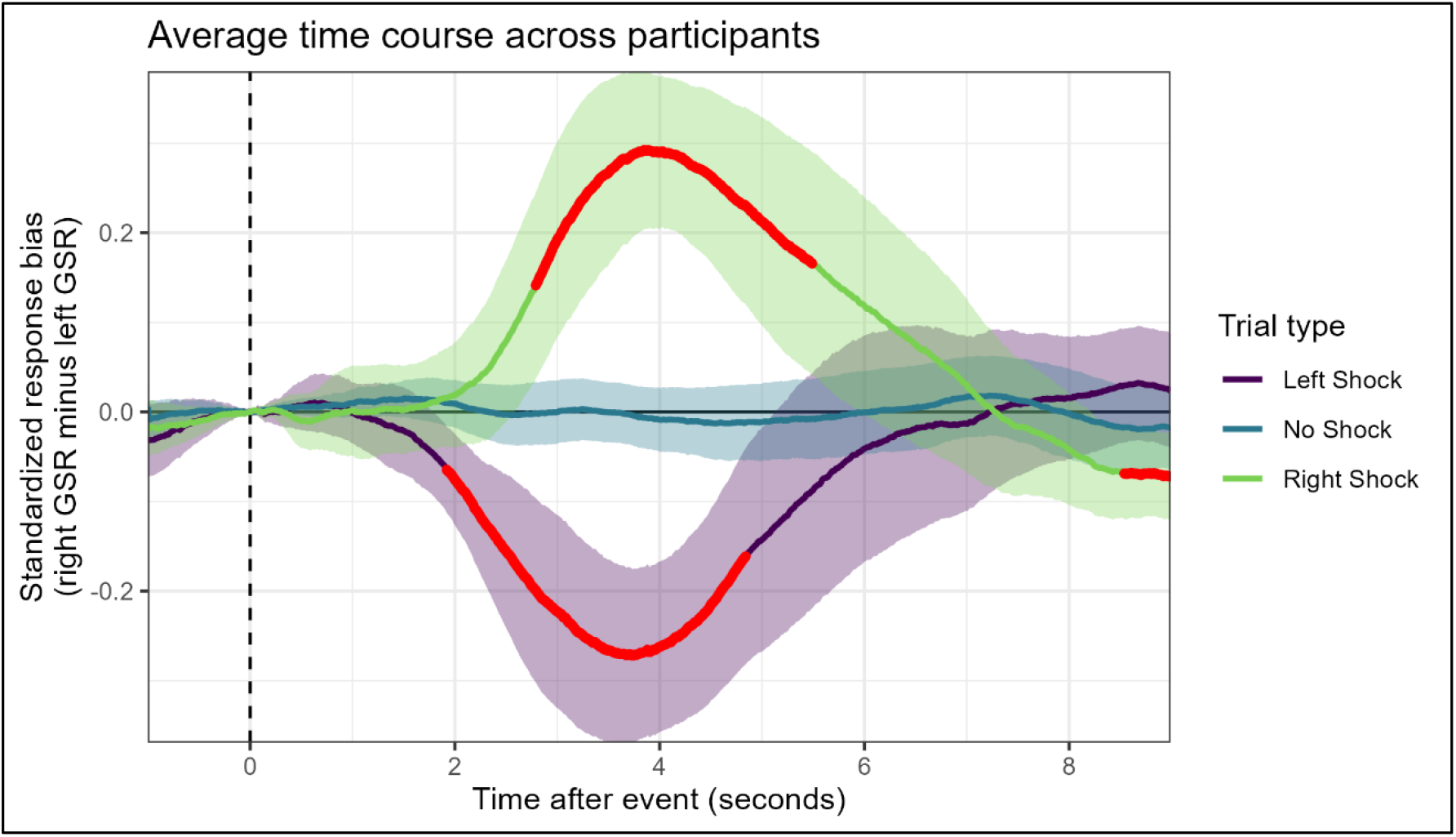
Group-wise contrast of lateralization bias in autonomic response. Lateralization Biases were defined as [Right-hand EDA – Left-hand EDA] for each condition (Left shock, Right Shock, and No Shock). Significant deviation of these Bias scores from a test value of zero (i.e., no bias) are indicated by the red highlight. Light colored ribbons represent the uncorrected 95% confidence interval at each time point.

### Within-subject analyses

If lateralized EDA to limb-specific threat is a foundational component of autonomic nervous architecture, this left vs right lateralization should be highly notable within individual subjects as well as across subject analyses. Accordingly, a similar set of analysis as presented for group-wise comparisons were conducted at a single subject level. For each subject, a series of one-sided t-test comparing Lateralization Bias for each individual trial type to null value of zero was conducted. Critically, these identified an overall pattern of results mirroring that observed at the group level (Figure 2a). During right-administered shock events, 26 of 50 participants displayed a lateralization bias in a direction that was consistent with those observed in the group-wise analyses. Similarly, during left-administered shock events, an overlapping (but not identical) group of 26 participants displayed lateralization bias in a consistent direction to the group-wise analyses. Of note, there were a maximum of 30 possible trials for each condition, with an average of 22.5 left shock events and 22.9 right shock events analyzed following data cleaning.

**Figure 2.**
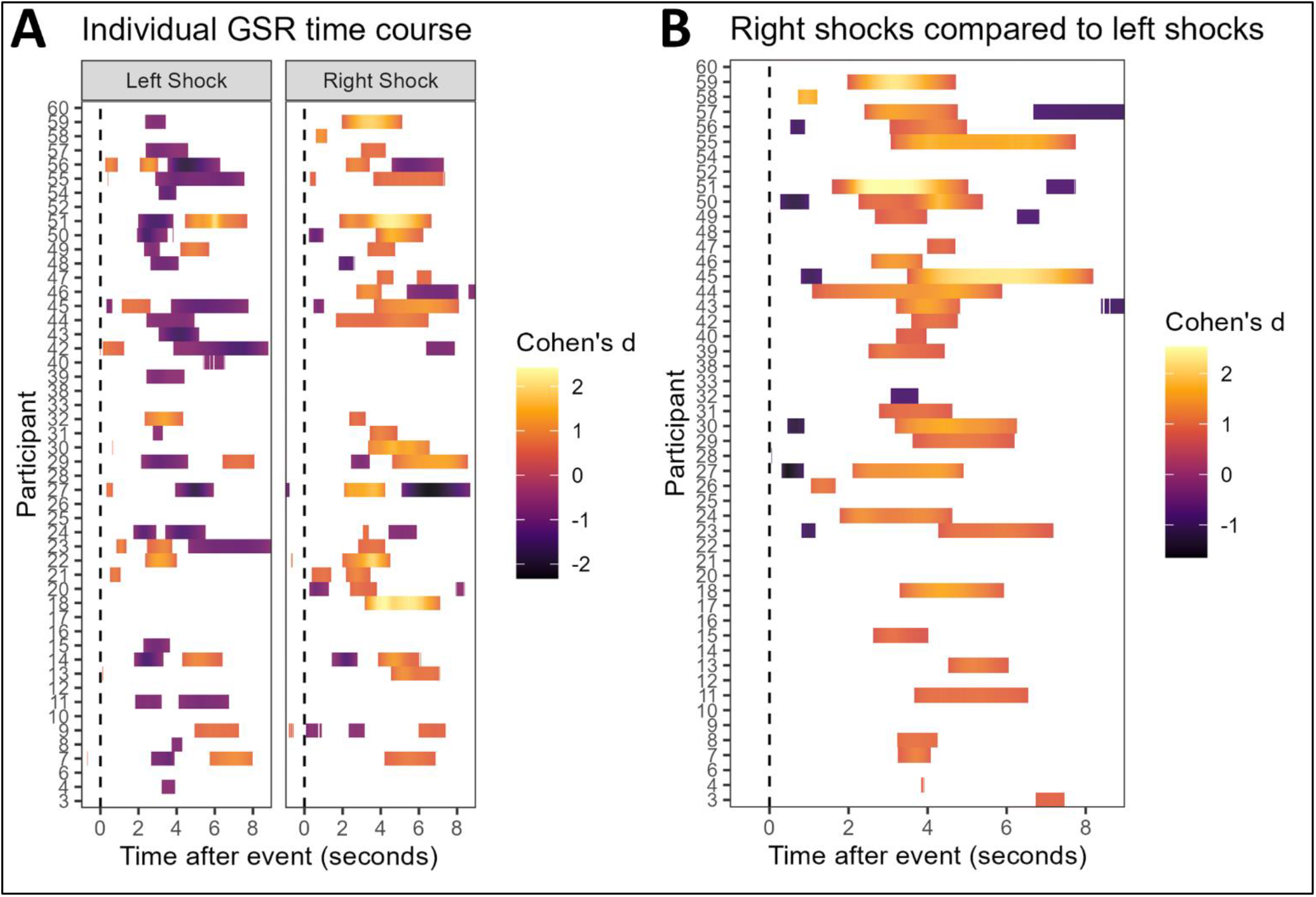
A. *Lateralization Biases across shock conditions by participant*. Differences in Lateralization Bias between left *versus* right shock events are identified for each participant (row) by trial time course. Cohen’s d is indicated for all time points where differences are significant at *p* < 0.05. B. *Lateralization Bias within shock conditions by participant*. Lateralization Bias scores for left *and* right shock events differing from no bias (i.e., Lateralization Bias = 0) are identified for each participant (row) by trial time course. Cohen’s d is presented for all time points where measured Lateralization Biases are significantly different from zero (*p* < 0.05).

To further reinforce the ubiquity of autonomic lateralization, an additional set of single subject analyses were performed on the trial-by-trial variability in Lateralization Bias for each participant. In a series of time-locked two-sample t-tests, significant differences in Lateralization Bias in EDA responses to shock administered to the left vs. right forearm were observed in 31 of 50 participants (Figure 2b). While the time course of significant differences ranged from as early as 500 ms post stimulus onset to as late as 8000 ms post stimulus onset, the most consistent range was 2000 – 5000 ms post onset, as was observed in the group analyses.

## Discussion

This study investigated whether lateralized electrodermal activity (EDA) would be observed in response to limb-specific tactile stimulation. For both left- and right-administered shock, EDA responses were larger at recording sites ipsilateral vs. contralateral to the stimulation site. This pattern of results was observed – quite strikingly – in group-wise analyses as well as at an individual subject level in more than half of all participants. Together, these findings provide strong evidence that the autonomic nervous system (ANS) exhibits robust specificity in EDA, which prioritizes responding in threat adjacent limbs.

While lateralized EDA in response to centrally-mediated, or general, states of arousal (e.g., faces, emotionally salient situations) has been periodically observed throughout the past century (for review of pre-1985 examples, see i Baque et al., 1984; also see: Bjorhei et al., 2019; Picard et al., 2016; Poh et al., 2010), a plausible functional rationale for this neuro-architectural quirk has remained elusive. Historical explanation often evoked now-discredited theories of gross hemispheric lateralization (e.g., Galaburda et al., 1978; Reeves, 1983; Sperry, 1968), while recent work has not addressed the role of asymmetric autonomic control in the context of metabolic conservation (Bjorhei et al., 2019; Picard et al., 2016). The current study demonstrates that lateralized EDA response is not only observed in response to centrally mediated arousal but can also be evoked by limb-specific aversive stimulation, thus providing a physiological rational for the development of asymmetric architecture in the ANS consistent with its canonical role of metabolic resource management. Specifically, increased EDA was observed in both the left and right arm following ipsilateral (vs. contralateral) electrical stimulation. This provides evidence for a functional role in threat response for lateralized changes in EDA. We suggest that the sympathetic output of our ANS – the driver of EDA (Vetrugno et al., 2003) – prepares the body for a response option beyond the popular alliterative of ‘fight or flight’; it can ready us to flick.

### Heterogeneity of autonomic output

Evidence of hemisphere-specificity in autonomic responses is consistent with the increasingly prevalent view that ANS output, and particularly ANS output in response to emotional arousal, should not be interpreted as a single measure of balance between global activation/conservation of resources (for review, see Kreibig, 2010). Further, this work demonstrates that heterogeneity of ANS outputs extends beyond differential signaling to separate effector organs, as it also includes differential signaling across body-locations within a single effector organ (i.e., skin). This is consistent with the view that, when a limb-specific motor response is required in response to threat, limb-specific ANS outputs increase local action-preparedness while a state of rest across the rest of the body is preserved. Through this mechanism, the global loss of metabolic resources can be minimized while still ensuring resources are available for adequate behavioural responses to threatening objects or situations (Gibbons, 2019; Wehrwein et al., 2016).

The current work is the first contemporary study to identify threat-localized lateralization in ANS responding and the first to observe it at a single subject level. As early preliminary work in the area (Fuhrer, 1971) was initially overlooked, and later disregarded entirely, much remains to be learned about the specificity of the autonomic response to localized threat. While we provide evidence for local specificity of cutaneous responses to tactile stimulation, it is unclear whether similar spatial heterogeneity also occurs in other ANS effector organs (e.g., vascular dilation), or in response to localized threats assessed by other modalities (e.g., visual threat). While some autonomic measures may best be regarded as homogenous metrics – e.g., cardiac and respiratory outputs – others are likely controlled in a heterogeneous manner to allow maximal metabolic conservation. For example, similar reasoning on the need to limit motor preparedness would suggest that vascular dilation should also be limited to threat targeted limb, rather than across the entire body. While the neural architecture for limb-specific vascular responding is well established – localized patterns of dilation are well documented during motor activity and exercise (Green, 2009; Green et al., 2017; Wang, 2005) – it is unclear whether this localized vascular responding can be used by the ANS for motor preparation as well. It is also unclear whether limb-specific cutaneous activity is observed in response to perception of threat through senses other than others, such the sight of a spider approaching the hand. Further investigation into modal specificity required to provoke body-localized changes in EDA would provide further insight into the functionality of such ANS responses, as well as highlight potential central structures involved in mediating these outputs.

### Development of cognitive arousal systems

A critical contribution of the current work is that we examined lateralized biases in electrodermal responding in response to direct physical threat to a distal *sensory target*, rather than manipulating or examining circumstances triggering emotional arousal to examine a centrally-mediated *cognitive state*. The importance of this difference becomes apparent in the context of understanding the biological role and evolutionary development of heterogeneity in cutaneous ANS functioning. To date, contemporary work in the field has largely ignored these questions, focusing instead on either the applied uses of asymmetric electrodermal signaling - including monitoring health and wellness (Arza et al., 2019; Carreiro et al., 2015a; Ghandeharioun et al., 2017; Greene et al., 2016; Kourtis et al., 2019) - or the cognitive states during which they manifest (Mohr et al., 2017; Picard et al., 2016). While these are important practical considerations when applying EDA signalling to healthcare concerns, neither provide any reasonable rationale for how the lateralized autonomic system they are describing may have developed.

By demonstrating greater EDA in body areas adjacent to vs. distant from tactile threat, we have provided evidence that supports the proposal that the lateralized neural architecture observed in the cutaneous ANS serves a concrete function in efficient threat protection. Furthermore, we propose that the observation of lateralized EDA in response to centrally mediated arousals (e.g., faces, Banks et al., 2012; high vs. low impact stressors, Bjorhei et al., 2019; situational arousal, Obrist, 1963; Picard et al., 2016) indicates that at some point during evolutionary development, central arousal systems likely co-opted pathways once dedicated to processing sensory arousal (Kryklywy et al., 2020). While the idea of building cognitive affect processing mechanisms on top of structures involved in sensory affect processing has been outlined for other specific sensory modalities (e.g., oral vs. moral disgust, Chapman et al., 2009) and as a modality general evolutionary practice (Kryklywy et al., 2020), these proposals often rely on the interpretation of shared patterns of activity within the brain, rather than shared peripheral outputs. The current work, however, highlights a specific example where the affective characteristic shared between general and tactile processes is peripheral in nature, yet only biologically sensible in its tactile manifestation. Additional work investigating the neural underpinning of the ANS modulation to both sensation provoked and centrally mediated arousals is still required to determine the extent to which these are overlapping processes within the central nervous system

## Methods

### Subjects

60 healthy participants (41F, Mean age = 20.45, SD = 2.6) were recruited from the UBC Psychology Human Subject Pool. Five of the subjects indicated they are left-handed. This study was approved by the Behavioural Research Ethics Board (BREB) at the University of British Columbia. Five participants were removed due to insufficient data or incomplete tasks (i.e., < 2 completed runs for each hand), and five more were removed due to excessive noise in the recording signal due to inadequate electrode contact or excessive participant motion (i.e., preprocessed data resulted in > 50% of individual trials discarded due to noise). Thus, all analyses described below were conducted on the remaining 50 subjects.

### Stimulus and Apparatus

All psychophysiological recording and stimulation were conducted through an AD Instrument Powerlab 8/35 DAQ device (PL3508). Tactile stimulation (Max repeat rate = 500Hz; and pulse width = 1ms; titrated voltage for each participant) was administered via stimulating bar electrodes (AD Instruments: MLADDF30) connected to an AD Instruments-Stimulus Isolator (FE180). Bar electrodes with a conductive gel were placed on bilaterally on participant forearms approximately 15cm distal to the elbow and fixed in place with medical tape. Pretrial stimulation was performed to ensure minimal activation of motor units in the arm and hand by the bar electrode. All stimulation events were 50 ms (pulse width = 1 ms, pulse height = 5 V, repeats = 50, repeat rate = 1000 Hz) and administered with a rectangular waveform. Strength of stimulation was titrated independently for each arm until participants reported that it was aversive but not painful. Titration involved increasing stimulus amplitude, beginning at 0.5 mA until a maximum of 10 mA until stimulation elicited a consistent self-report rating of aversive but not painful. Increments varied from 1 mA to 0.1mA depending on the previous response to stimulation.

EDA was collected bilaterally from by galvanic skin response (GSR) finger electrodes (AD Instruments: MLT118F) placed on the middle and index fingers of each hand and amplified by a GSR-amp (AD Instruments: FE116). In addition, electrocardiogram (ECG) was collected with a single finger pulse transducer (AD Instruments: TN1012/ST) connected to the left thumb. An attentional visual task is run on PsychoPy v1.90.1, presented on a monitor (BenQ XL Series XL2420B 24 Widescreen; 60hz) placed ∼60cm away from the participant. Initial EDA preprocessing was conducted with Labchart 8 (AD Instruments). All data was sampled at 1000Hz.

### Procedure

Prior to data collection, participants were asked to sit upright and in a comfortable position, with their hands placed on a table, to minimize movement throughout the experiment. Participants completed six blocks of a task designed to monitor bilaterally EDA response to aversive tactile stimulation. Over the duration of each block, participants received ten electrical shocks to one arm. The target of stimulation (left vs. right forearm) alternated between blocks, with the initial location counterbalanced across participants. Shock events were spaced 10,000 ms apart and intermixed with an equivalent number of ‘No Shock’ trials. Participants were instructed to remain still for the duration of the experiment and told that no response to the tactile stimuli was required at any point during the task. To minimize participant motion, and maintain engagement, participants were instructed to track the number of colour changes in a centrally-presented fixation cross and made a verbal report of this count to the experimenter following each block (range = 40-60 changes per block). Over the duration of the experiment, participants received a maximum of 60 shock events (30 to each arm), as well as 60 ‘No Shock’ events. Bilateral EDA was collected over the duration of all events.

### Preprocessing and Analyse

EDA data was exported from Labchart and down-sampled to 100 Hz (from 1000 Hz) to facilitate subsequent analyses. EDA from the electrode on each hand was subjected to second order Butterworth filter with a bandpass of 0.05 to 30 Hz. An additional manual filter of unlabeled trial-by-trial data was conducted to identify a GSR threshold beyond which the data was most likely attributed to noise in the signal. GSR thresholds were identified independently by two unique raters. Inter-class correlation estimates found excellent inter-rater reliability (Cronbach’s alpha = 0.891), and so thresholds identified by Rater 1 were used for all remaining analyses. Remaining data was subject to z-score standardization for each signal (i.e., by GSR electrode). All shock events were extracted to produce a lateralized bias score. For each independent event (i.e., all shock and no shock events), we calculated the right hand GSR – left hand GSR for each time point, resulting in a continuous ‘Lateralization Bias’ index for each event, with positive values indicating right-biased asymmetry, and negative values indicating left-biased asymmetry. For group analyses, data from each participant was collapsed across trial type, resulting in three unique conditions (‘Left Shock’, ‘Right Shock’, and ‘No Shock’). Three series of one-sample t-tests were conducted comparing the Lateralization Bias for each condition against a test value of 0 (i.e., no bias) at each time point, with all resultant *p* values subject to a false discovery rate correction (Benjamini and Hochberg, 1995). For single-subject analyses, trial events remained separate, and were used as independent repetitions in subsequent inferential tests. A series of two-sample t-tests compared lateralization biases during left and right shock events to identify whether the group effects were observable within single individuals, in addition to one-sample t-tests for both right and left shock conditions. As with group analyses, all resultant *p* values were subject to a false discovery rate correction.

